# PP-MAPS: dynamic pharmacophore signatures of protein–peptide interfaces from molecular dynamics trajectories

**DOI:** 10.64898/2026.04.22.720140

**Authors:** Camille Depenveiller, Arezki Guerda, Emilia Rabia, Aziza Caidi, Yaqoub Ashhab, Fathia Mami-Chouaib, Matthieu Montes

**Author notes:** Corresponding authors (Camille Depenveiller; Matthieu Montes).

## Abstract

Protein–peptide interactions underlie many cellular signaling and regulatory processes and are increasingly exploited in drug discovery. Characterizing such interfaces often requires the analysis of ensembles of conformations obtained by molecular modeling or molecular dynamics (MD) simulations, where transient contacts and alternative binding modes can be critical. Pharmacophore models provide an intuitive, transferable representation of molecular interactions. Dynophore i.e. “dynamic pharmacophore” approaches have been developed for small-molecule ligands with MD information. We present PP-MAPS (Protein-Peptide Molecular dynamics Assisted Pharmacophore Signatures), an open-source workflow that extracts and aggregates pharmacophore interactions along MD trajectories of protein–peptide complexes. PP-MAPS produces per-residue interaction frequencies and pharmacophore heatmaps that facilitate comparison of peptides, binding sites and receptor variants.

PP-MAPS is implemented in Python and is available under an open-source license at https://github.com/camilledepenveiller/PP-MAPS. The workflow relies on GROMACS for trajectory processing and can use either LigandScout or the Chemical Data Processing Toolkit (CDPKit) for pharmacophore feature detection.

## Introduction

Protein–peptide interactions are pervasive in biology, ranging from short linear motif recognition to high-affinity complexes, and they contribute substantially to cellular regulation and disease mechanisms^1,2^. Because peptides can achieve high specificity with relatively low toxicity, they are also an expanding class of therapeutics^3,4^. A major challenge is that peptide binding interfaces are often shallow and dynamic, with induced fit and multiple metastable interaction patterns. Molecular dynamics simulations provide avenues to sample this conformational heterogeneity at atomic resolution^5^, but extracting actionable, comparable interaction information from trajectories remains non-trivial.

Recent advances in structure prediction and docking, ranging from flexible protein–peptide docking frameworks^6,7,8,9,10^ to deep generative approaches for molecular complexes^11,12,13,14,15,16,17,18^, have enabled the rapid proposal of protein–peptide binding modes. These developments also support computational design and optimization of peptide-based ligands^19^. Downstream analysis tools are still needed to interpret and compare predicted or simulated conformational ensembles, especially when the goal is to optimize peptide sequences for binding or understand key binding determinants.

Pharmacophore models encode the spatial arrangements of interaction features (e.g. hydrogen-bond donors/acceptors, hydrophobic, aromatics and charged groups) that ensure optimal interactions with the target and are widely used in ligand-based and structure-based drug design^20,21^. Pharmacophores can also be represented and compared using graph-based encodings, facilitating transferable interaction signatures^22^. Several methods extend pharmacophores to MD by computing “dynamic pharmacophores” or “dynophores” for small-molecule ligands^23,24,25,26^. However, peptides can pose practical difficulties: since they can be encoded as part of the macromolecule, the ligand/receptor partition is not always explicit in simulation outputs. PP-MAPS addresses this gap by providing a streamlined workflow to compute and summarize pharmacophore interactions for protein-peptide complexes along MD trajectories.

As an illustrative use case, we applied PP-MAPS to different peptide–major histocompatibility complex (pMHC) complexes. pMHC interactions determine antigen presentation and T-cell recognition, underpinning cancer immunotherapy and vaccine development^27,28^. While affinity-based predictors such as NetMHCpan^29,30^, MixMHCpred^31^, MHCflurry^32^, DeepAttentionPan^33^ and CapsNet-MHC^34^ provide valuable peptide selection, complementary approaches based on receptor pocket similarity have also been proposed^35^, as well as pMHC model predictors^36,37^. Structural and dynamic analyses can further reveal how different peptides exploit MHC pockets and which contacts are stable or transient i.e an information that PP-MAPS captures in a compact and standardized representation.

### Implementation

PP-MAPS takes as input a topology file (GROMACS TPR) and a trajectory (GROMACS XTC) of a protein–peptide complex. The workflow uses GROMACS^38^ to extract individual frames in the PDB format and enforces an explicit protein/peptide partition with the peptide placed at the end of each PDB file. This step ensures compatibility with pharmacophore engines that expect a macromolecular receptor and a distinct ligand.

For each frame, pharmacophore features and interactions are detected using either LigandScout^20^ or the open-source CDPKit backend^39^. Detected interactions are assigned to receptor residues and classified by type (hydrogen bond, ionic, hydrophobic, aromatic, etc.). All frame-level interaction sets are stored in a structured JSON output.

PP-MAPS then aggregates interactions across the trajectory to compute frequencies and produce pharmacophore “heatmaps”: a matrix of interaction types versus receptor residues, where values correspond to occurrence frequencies along the MD trajectory i.e the conformational ensemble. This allows to further perform per-residue contact statistics, extraction of high-frequency pharmacophore features, and 3D visualization to project frequent features onto a reference structure.

### Use case: pharmacophore signatures of peptide–MHC complexes

To demonstrate the use of PP-MAPS on a biologically and therapeutically relevant family of protein–peptide interactions, we analyzed three MHC of type HLA-A*02:01 complexes presenting T-cell epitopes associated with impaired peptide processing (TEIPP) peptides, which can be displayed independently of the transporter associated with antigen processing (TAP) and are of interest for targeting immune-escaped tumors^40,41^. Starting from the template structure (PDB 6TRN), we modelled three peptides: A: VLLQAGSLHA^42^, B: MLLAVLYCL^43^ and C: LSEKLERI^44^. pMHC complexes were generated with recent structure prediction and complex modelling frameworks^11,37,16,14,17^.

All complexes were simulated with GROMACS^38^ using identical parameters (500 ns production run; 2 fs time step; explicit solvent; 0.15 M NaCl). From each trajectory, 50,000 conformations were extracted and processed by PP-MAPS. The following command line was used for each complex:

**python pp_maps.py -xtc md.xtc -tpr md.tpr -ligandscout -n 8**, with md.xtc and md.tpr the input files from MD, LigandScout as chosen software and n the number of processes. The output file is a PNG file of the pharmacophore heatmap.

The resulting pharmacophore heatmaps presented in Figure 1 highlight both conserved and peptide-specific interaction patterns, enabling rapid comparison of how peptides occupy MHC pockets and interact with key residues.

**Figure 1.**
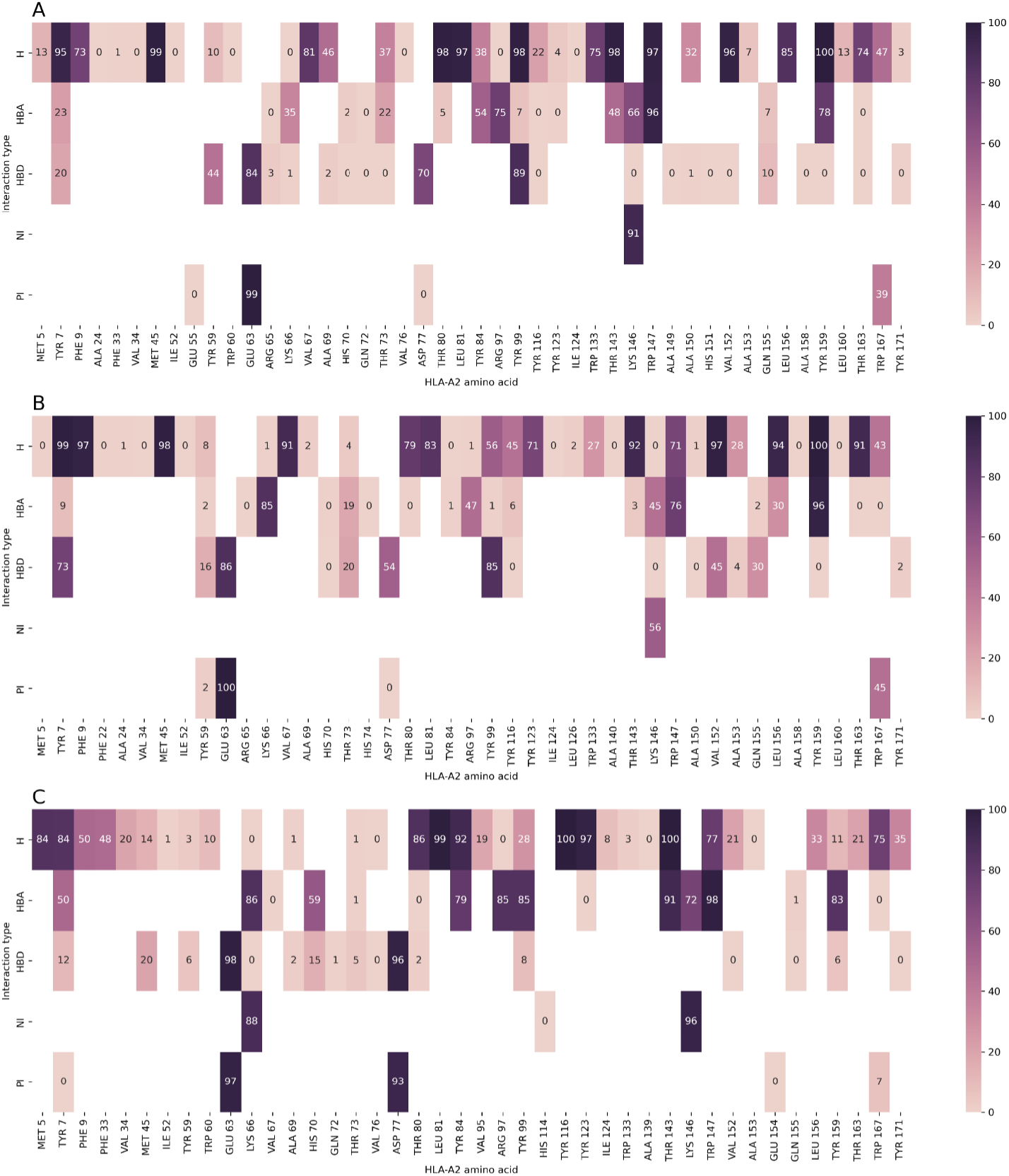
Pharmacophore heatmaps produced by PP-MAPS for three TEIPP peptides (A: VLLQAGSLHA; B: MLLAVLYCL and C: LSEKLERI) bound to HLA-A*02:01. Rows correspond to pharmacophore interaction types and columns to MHC residues; values denote the frequency of each interaction along the MD trajectory from light pink (0%) to dark blue (100%). Shared high-frequency contacts are readily apparent across peptides, while peptide-specific signatures reveal alternative pocket usage.

Across the three complexes, PP-MAPS consistently recovered a core set of common pharmacophore signatures between peptides, corresponding to redundant frequent interactions involving the A*02:01 binding groove. These pharmacophore signatures included hydrophobic interaction with TYR 7, H-bond donor and positive ionizable with GLU 63, hydrophobic with THR 143, negative ionizable with LYS 146, hydrophobic and H-bond acceptor with TRP 147, H-bond acceptor with TYR 159. The corresponding amino acids were evidenced on the structure of the complexes.

Figure 2 shows the structural positioning of the signature pharmacophore interactions found for the VLLQAGSLHA-MHC complex. These contacts were established around the known anchor positions of TEIPP^42^, evidenced in this analysis with PP-MAPS. Differences between peptides were captured as distinct pharmacophore signatures that can be used to 1. compare candidate epitopes presenting similar predicted affinities but different structural stabilization strategies, and 2. identify peptide positions that could be modified without disrupting anchor-like interactions.

**Figure 2.**
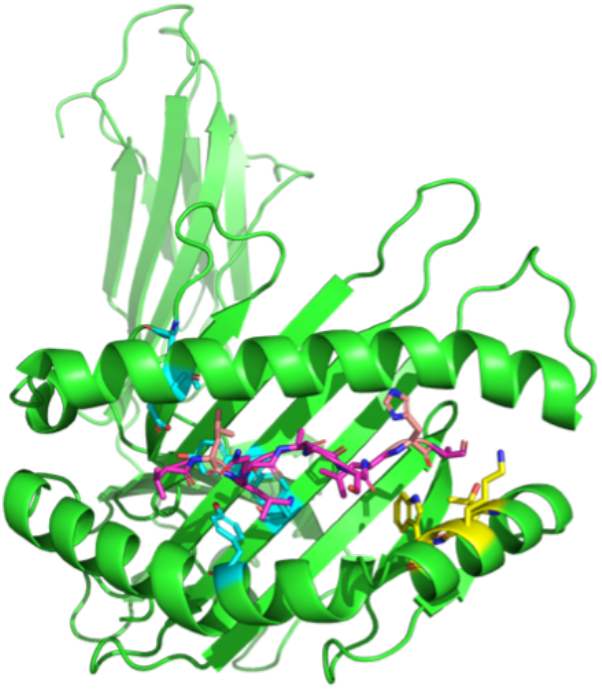
VLLQAGSLHA epitope in complex with HLA-A*02:01 and interacting residues. HLA-A*02:01 is colored in green and VLLQAGSLHA in magenta, with anchor positions 2 and 9 in light pink. HLA-A*02:01 amino acids that were evidenced by PP-MAPS as the most involved in interactions are colored in cyan when surrounding VLLQAGSLHA position 2 and in yellow when surrounding position 9.

In practical epitope discovery pipelines, PP-MAPS can be used downstream of binding predictors (e.g. NetMHCpan/MixMHCpred/MHCflurry/DeepAttentionPan/CapsNet-MHC) and structure modeling to prioritize peptides whose trajectories maintain stable, well-distributed pharmacophore contacts, or to understand why a peptide with a favorable binding score yields weak or unstable presentation.

## Conclusion

PP-MAPS provides an automated, open-source approach to summarize protein–peptide interactions along MD trajectories using pharmacophore representations. By converting conformational ensembles into compact pharmacophore signatures, PP-MAPS facilitates comparison across peptides and receptor variants and supports data-driven hypothesis generation for peptide optimization. The pMHC example illustrates how pharmacophore heatmaps can complement affinity prediction by revealing dynamic contact patterns within the MHC groove. More generally, the same workflow applies to any protein-peptide system where conformational ensembles are available (from MD, docking refinement or alternative structural models), providing a comparable representation for peptide optimization and interface engineering.

## Data and software availability

PP-MAPS is available under an open-source license, all scripts and files are provided at https://github.com/camilledepenveiller/PP-MAPS.

## Author Contributions

Camille Depenveiller: Investigation, Software implementation, Analysis, Writing - original draft; Arezki Guerda: Analysis; Emilia Rabia: Investigation, Analysis; Aziza Caidi: Analysis; Yaqoub Ashhab: Investigation, Analysis; Fathia Mami-Chouaib: Investigation, Analysis, Writing - review; Matthieu Montes: Conception, Investigation, Analysis, Writing – review.

## Acknowledgments

We thank Thierry Langer and Inte:ligand for kindly providing a LigandScout license.

## Abbreviations

MD: molecular dynamics
pMHC: peptide-major histocompatibility complex
TAP: transporter associated with antigen processing
TEIPP: T-cell epitopes associated with impaired peptide processing.

## Graphical Abstract

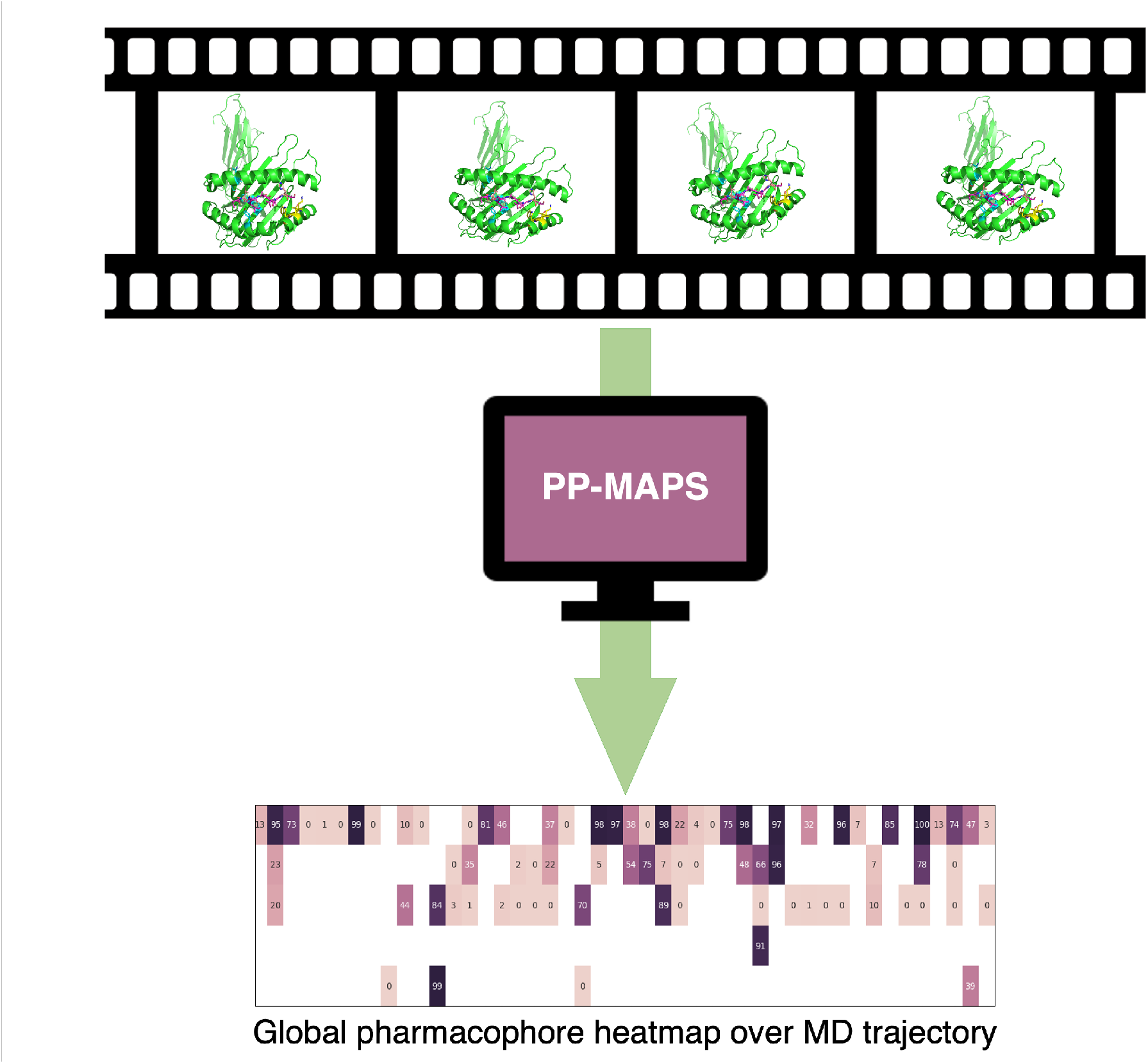

